# Development and design rules for a modular human peptide-tag adaptor CAR T cell system

**DOI:** 10.64898/2026.10.21.760864

**Authors:** Alexander Haide Joechner, Melanie Mach, Michael Yeh, Riya Mehta, Taliah Khoury, Ecem Kurangil, Jinting Yan, Daniel Barton, Ziduo Li

## Abstract

CAR T cells are highly effective in haematological malignancies, but limited post-infusion control and tumour antigen heterogeneity remain major challenges. Modular CAR systems address these limitations using interchangeable adaptor molecules (AM) to control CAR T cell engagement and redirect targeting toward different antigens, yet the design principles underlying effective modular function remain undefined. We present our modular platform, BRiDGE CAR, and systematically examine how CAR and adaptor configuration influence activity. BRiDGE CAR activity was influenced by CAR hinge length and adaptor format and valency; a medium-length hinge combined with a full-length, bivalent adaptor supported potent activity across multiple tumour antigens and consistently controlled tumour growth across the tested xenograft models. During repeated antigen challenge, CD33-directed BRiDGE CAR T cells retained less-differentiated phenotypes than direct CD33-targeting CAR T cells. *In vivo*, BRiDGE CAR T cells persisted during adaptor withdrawal and regained antitumour activity upon adaptor re-administration, while IL-18 armouring enhanced tumour control at a reduced CAR T-cell dose. We identified key design parameters governing rules for adaptor-mediated CAR activity, with antigen targeting and CAR T cell function remaining independently modifiable, establishing BRiDGE CAR as a flexible and controllable modular platform.

## Introduction

CAR T cell therapy achieves deep and durable remissions in B-lineage haematological malignancies and multiple myeloma (MM) [1–3] but its success has not extended to acute myeloid leukaemia (AML) or most solid tumours [4–6]. In AML, a major challenge is the lack of suitable tumour-associated antigens (TAA), since many candidate targets are also expressed on normal haematopoietic cells, creating a risk of on-target/off-tumour (OTOT) toxicity [7–10]. In solid tumours, heterogeneous antigen expression can limit tumour coverage and promote antigen escape, while the tumour microenvironment further restricts CAR T cell function and persistence [11, 12]. In conventional CARs, the target antigen is also built into the receptor, so it cannot be changed or switched off once the cells are given.

Modular CAR systems offer a solution by separating antigen recognition from CAR signalling. A soluble adaptor molecule (AM) binds the tumour antigen and carries a tag recognised by a universal CAR, thereby bridging the CAR T cell to its target [13–19]. This architecture allows CAR activity to be regulated through adaptor availability and enables the same universal CAR T cells to be redirected by exchanging or combining AMs. However, introducing an intermediate recognition molecule creates additional engineering constraints. CAR–AM–antigen geometry, adaptor format and valency can influence activity, while adaptor pharmacokinetics affect the duration and reversibility of CAR T cell activation. Several adaptor CAR platforms have also been reported to show lower antitumour potency than their conventional CAR counterparts [20, 21].

A central challenge in engineering modular CAR platforms is therefore to define design rules that achieve antitumour effectiveness without losing modular flexibility. Open questions include how the CAR and adaptor should be configured, how adaptor concentration and target antigen density together set CAR T cell activity, and whether the resulting rules hold across different tumour antigens. Defining these relationships is important for developing modular CAR systems that combine robust anti-tumour activity with control over antigen recognition.

We developed the BRiDGE CAR, a modular system based on the E200 peptide epitope derived from the non-functional conformation of human P2X7 (nfP2X7) and its cognate antibody Bil03 [22]. We systematically examined the design principles governing modular CAR activity and tested their applicability across multiple haematological and solid tumour models. We also compared the response of adaptor-engaged BRiDGE CAR T cells with direct CAR T cells during repeated antigen challenge and explored the versatility of the platform through adaptor re-dosing, bispecific targeting and armouring CAR T cells with inducible IL-18 secretion.

## Results

### Hinge length and adaptor format determine BRiDGE CAR function

Unlike a conventional CAR, which recognises a tumour antigen directly, an adaptor CAR engages its target through a separate AM. We designed the BRiDGE system using an AM that binds a tumour antigen and carries a short peptide tag recognised by the CAR T cell, thereby bridging the two cells (Figure 1A). The BRiDGE CAR recognition epitope and paratope originate from human nfP2X7 [22, 23] and incorporate the E200 tag, a 17-amino-acid (aa) peptide and its corresponding antibody Bil03, from which the CAR binding domain is derived. Bil03 bound the E200 tag with a dissociation constant (K_D_) of 10.4 nM and was characterised by rapid association and dissociation kinetics (k_on_ = 7.288 × 10□M□¹ s□¹, k_off_ = 7.61 × 10□³ s□¹). The BRiDGE CAR is a third-generation receptor containing the Bil03 binding domain, hinge and transmembrane domains, and the intracellular costimulatory domains, CD28 and 4-1BB (Figure 1B).

**Figure 1.**
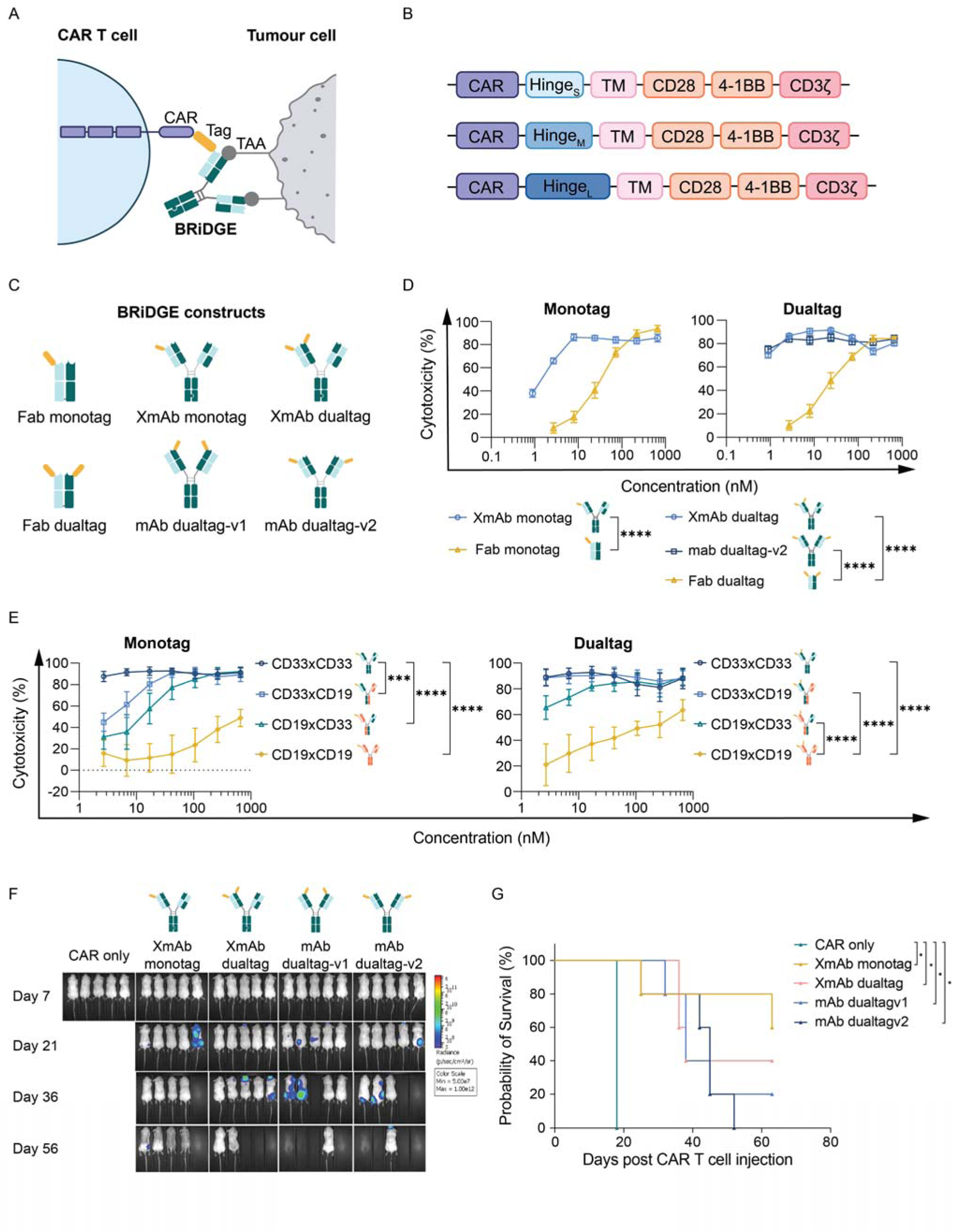
BRiDGE CAR design and development. **A** BRiDGE CAR mode of action: TAAs on the tumour cell are recognised by the AM (BRiDGE) and the CAR T cell recognises the AM via the tag. **B** BRiDGE CAR schematic with binding domain (CAR; Bil03-derived VH binder), a short, medium or long hinge, transmembrane (TM) domain as well as two costimulatory domains (CD28, 4-1BB) and the signalling domain (CD3ζ). **C** AM formats: single- or dual-E200 tag AMs were generated using different antibody scaffolds, including Fab, asymmetrical CrossMab antibody (XmAb) and conventional symmetric antibody (mAb) formats. **D** Luciferase assay comparing Fab and full-length antibodies against U937 in monotag (left; *P*<0.0001) and dualtag (right; *P*<0.0001 Fab vs XmAb and Fab vs mAb dualtag-v2) configuration (N=3 donors). Data presented as mean with SD, *P*-values calculated by two-way ANOVA with Tukey’s multiple comparisons test. **E** *In vitro* comparison of CD33×CD19 bispecific AM with CD33 or CD19 bivalent molecules comprising monotag (left) and dualtag (right) configurations in CD33^+^CD19− U937 cells (N*=*3 donors). Data presented as mean with SD, *P*-values calculated by two-way ANOVA with Tukey’s multiple comparisons test. **F** Bioluminescence imaging of all tumour-bearing mice treated with BRiDGE CAR T cells alone or with CD33 XmAb and mAb AM in mono- and dualtag configuration (N=5 mice per group). **G** Kaplan–Meier survival of the mice shown in F, *P*-values calculated by log-rank (Mantel–Cox) with multiple comparisons test CAR T only vs XmAb monotag *P*=0.0108, all others not significant (ns).

Because hinge length determines the spacing between the CAR T cell and tumour cell, we compared short, medium and long hinges for CAR expression, T cell expansion and *in vitro* cytotoxicity. Short and medium hinges showed comparable CAR expression and expansion, whereas the long hinge showed higher surface CAR expression but poorer expansion (Supplementary Figure S1A). Across three target antigens (CD19, CD33 and CD276/B7-H3), the medium hinge was the only configuration that retained cytotoxic activity in every model (Supplementary Figure S1B) and was therefore selected for all subsequent experiments.

Because the AM can be built on different antibody scaffolds, we examined how scaffold format and E200 tag valency affect BRiDGE CAR activity. We generated AMs based on Fab, conventional mAb and heterodimeric CrossMab (XmAb) scaffolds, carrying one E200 tag (monotag) or two E200 tags (dualtag) positioned on the same or opposite arms where applicable (Figure 1C) [24]. At equivalent molar concentrations, full-length antibody formats were substantially more potent than Fab-based AMs, with an approximately 1.5–2-log shift in EC_50_ (Figure 1D). In contrast, differences between mono- and dualtag formats within the antibody framework were modest, indicating that scaffold format had a greater influence on *in vitro* potency than tag valency.

To distinguish the contribution of binding valency from that of the scaffold, we tested CD33×CD19 bispecific AMs against CD33^+^CD19^−^ U937 cells, thereby restricting target engagement to a single CD33 binding arm. Bivalent CD33×CD33 AMs showed greater potency than the monovalent bispecific formats (Figure 1E), which retained greater activity than the Fab AM (Figure 1D). These findings indicate contributions from both bivalent target engagement and the full-length antibody format. Tag position also influenced activity: CD33×CD19, with the E200 tag on the target-engaging CD33 arm, was more potent than the reciprocal CD19×CD33 format, in which the tag was positioned on the non-engaging arm (Figure 1E).

We next compared CD33 AM formats *in vivo* in a MOLM-13 xenograft model to assess the impact of tag valency directly. The XmAb monotag AM provided the most durable tumour control and survival, whereas dualtag formats showed earlier relapse, with XmAb dualtag performing slightly better than mAb dualtag (Figure 1F, G). Consistent with this finding, a Trop2 XmAb monotag AM led to better *in vivo* tumour control than the corresponding dualtag format in an OVCAR-3 xenograft model (Supplementary Figure S1C, D).

Together, these studies show that hinge length, antibody format and E200 tag valency collectively determine BRiDGE CAR performance. Based on these findings, a medium-length hinge CAR and bivalent XmAb adaptor carrying a single E200 tag were selected for subsequent experiments.

### BRiDGE CAR activation requires recognition, integrity, and engagement of tag and TAA

The BRiDGE CAR target recognition requires coordinated interactions between the CAR, AM and TAA. To establish the contribution of each component, we independently disrupted tag recognition, tag integrity and antigen engagement (Figure 2A). Mutation of the CDR3 regions of the CAR binding domain abolished cytotoxicity against CD33^+^ U937 cells, confirming that CAR recognition of the E200 tag is required for activation (Figure 2B). Next, we systematically altered the E200 tag sequence, replacing each position with alanine and assessing the impact on affinity, cell-surface staining and cytotoxicity. Changes in affinity were reflected in surface staining and cytotoxicity, with five residues (I208, L209, P210, L212 and I214) consistently identified as critical across all three assays (Figure 2C). These residues therefore define the functional E200 epitope required for BRiDGE CAR activation. Lastly, to examine antigen specificity, we created CD33 and/or CLL-1 knockout variants of U937 cells, which were co-cultured at equal ratios with WT cells. BRiDGE CAR T combined with CD33 targeting AM selectively depleted the CD33^+^ subpopulation, whereas CLL-1 targeting depleted the CLL-1^+^ subpopulation. A CD33×CLL-1 bispecific AM eliminated cells expressing either antigen (Figure 2D). This finding was also confirmed using CD19 and CD20 AMs against WT and CD19-KO JeKo-1 cells (Supplementary Figure S2A, B).

**Figure 2.**
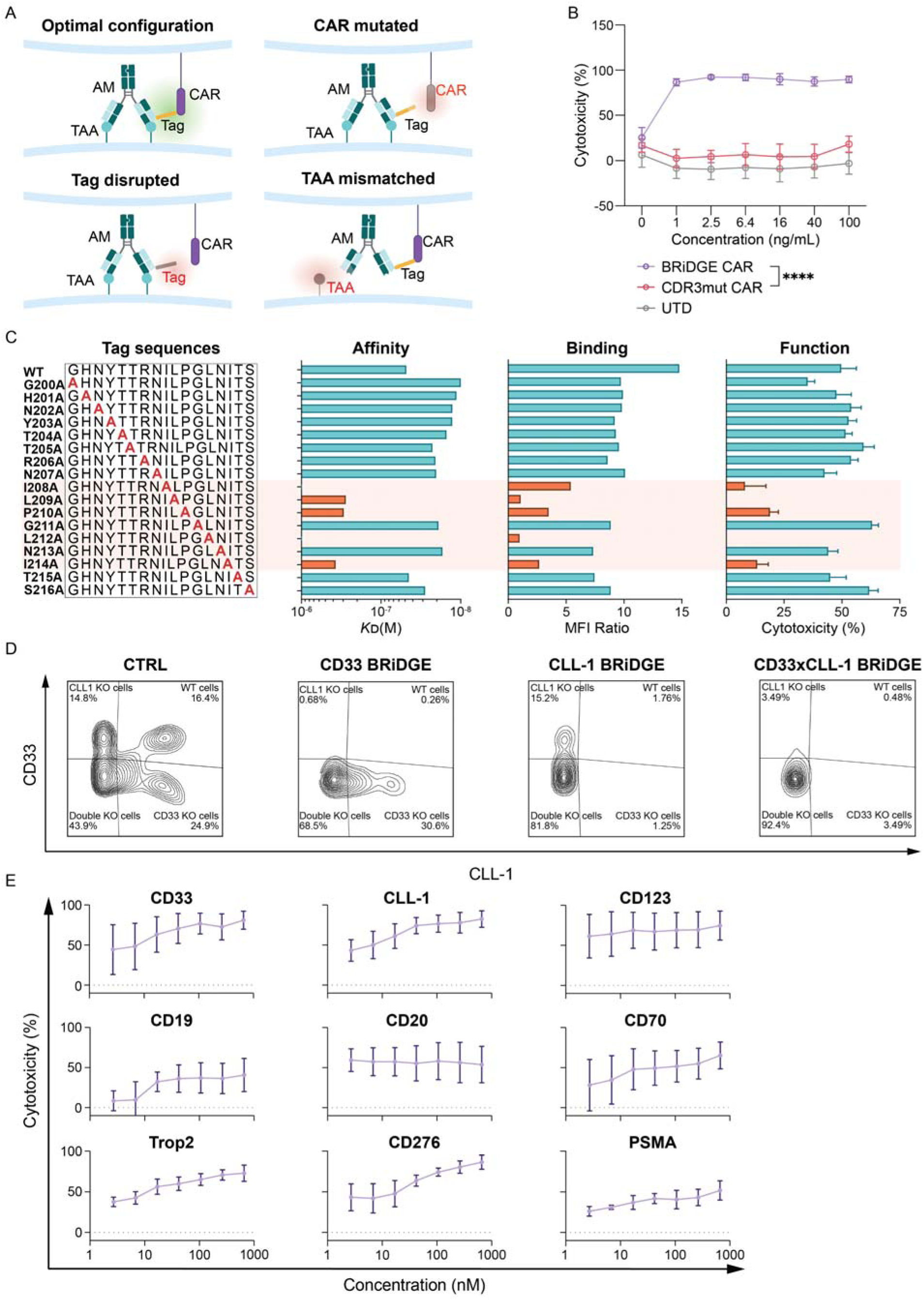
BRiDGE CAR activation requirements, specificity and versatility. **A** Schematic of the requirements for BRiDGE CAR activation: a correctly assembled CAR–AM–TAA chain leads to activation (top left), whereas a mutated CAR abolishes tag binding (top right), a disrupted tag abolishes CAR binding (bottom left), and a mismatched TAA abolishes AM binding to the target cell (bottom right). **B** Cytotoxicity data of U937 cells treated with a CDR3 mutated BRiDGE CAR compared to standard BRiDGE CAR and UTD T cells. N=3 donors, data presented as mean with SD, *P*-values calculated by two-way ANOVA with Tukey’s multiple comparisons test (*P*<0.0001 CAR vs CDR3mut and CAR vs UTD). **C** Systematic replacement of amino acids in our tag sequence with alanine (far left) resulted in AMs with reduced affinity (middle left), fluorescence intensity (middle right) and cytotoxicity (far right). Highlighted in red are the residues critical for function (I208A, L209A, P210A, L212A and I214A). L212A was below the limit of detection in all three assays, indicating complete abrogation. Representative plots for KD and MFI data, cytotoxicity data generated from N=2 donors, data presented as mean with SD. **D** Representative flow cytometric plots of CD33 and CLL-1 positive U937 cell subpopulations after 24 h co-incubation with BRiDGE CAR and CD33/CLL-1 mono- and bispecific AMs. **E** Cytotoxicity analysis of nine TAA targeted by BRiDGE CAR and indicated AMs, N=3 donors, data presented as mean with SD.

We investigated the versatility of the BRiDGE CAR by pairing the same BRiDGE CAR construct with AMs against nine antigens across haematological and solid tumours. Each AM bound its respective target cells (Supplementary Figure S2C) and mediated cytotoxicity against the corresponding tumour cells (Figure 2E), demonstrating that the same CAR construct can be applied to target different antigens by exchanging the AM.

### Adaptor-dependent activation and memory phenotype retention in BRiDGE CAR T cells

The optimised BRiDGE CAR and CD33 AM pair was characterised extensively in the AML setting. Titration of the CD33 AM across a panel of CD33^+^ AML cell lines resulted in dose-dependent cytotoxicity, with minimal activity against CD33^−^ JeKo-1 cells (Figure 3A). More extensive titration experiments of AM dose and E/T ratio showed that cytotoxicity depended on both parameters, with maximal killing at E/T ≥ 1 and AM concentrations ≥ 1 ng/mL (≈6.7 pM) (Supplementary Figure S3A). Similarly, IFN-γ and IL-2 secretion required the presence of both BRiDGE CAR T cells and AM in a dose-dependent manner (Figure 3B).

**Figure 3.**
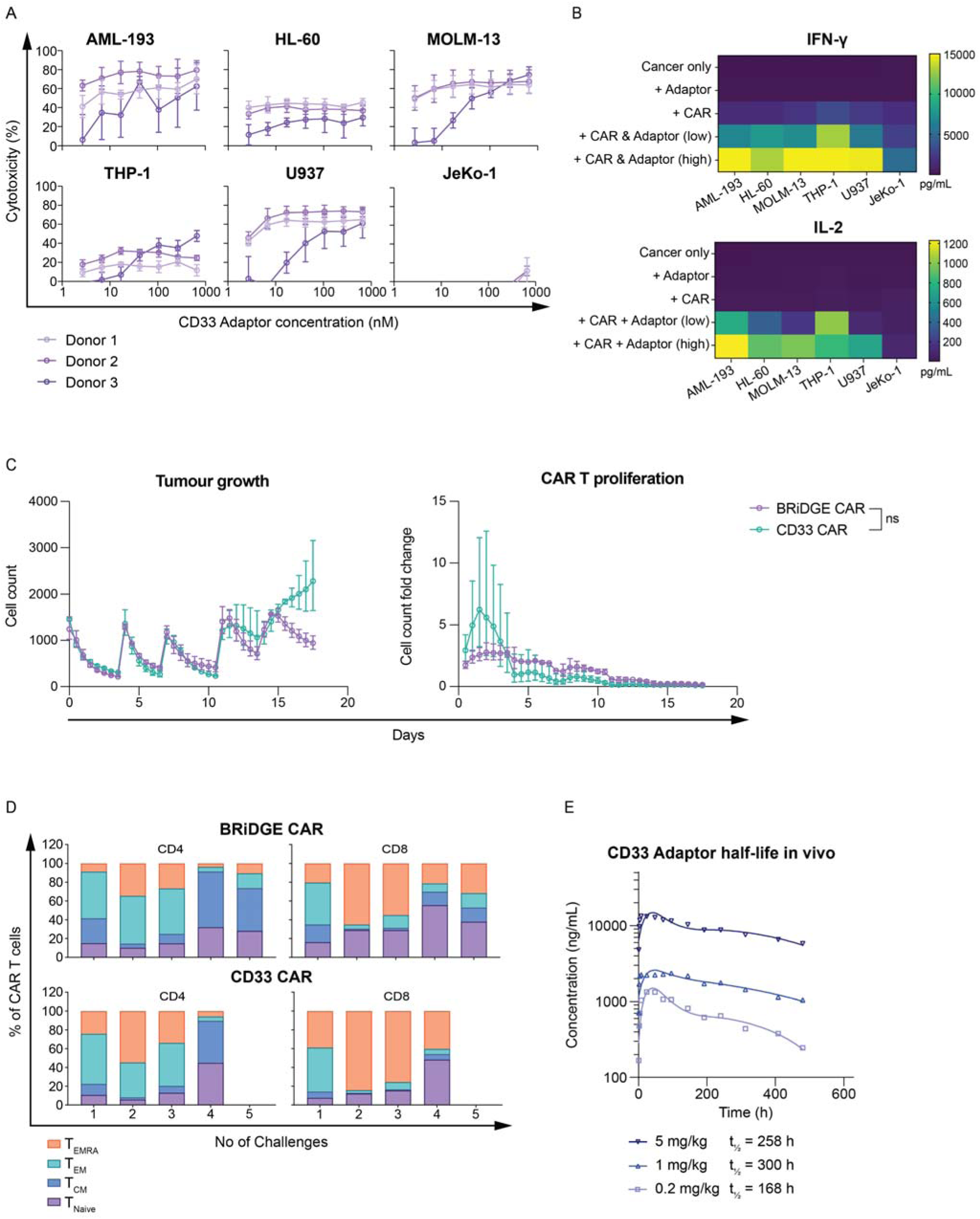
*In vitro* BRiDGE CAR efficacy in an AML setting and *in vivo* half-life analysis of AM. **A** Cytotoxicity of CD33 redirected BRiDGE CAR T cells vs CD33^+^ AML cell lines and CD33− JeKo-1 as control. N=3 donors, data presented as mean with SD. **B** Analysis of IFN-γ and IL-2 secretion levels in supernatants harvested from A. Conditions analysed were: cancer cells only, cancer cells treated with AM only, cancer cells treated with BRiDGE CAR T only, and cancer cells treated with BRiDGE CAR T cells plus AM. Two concentrations of CD33 AM shown: low = 42.7 pM (6.4 ng/mL) and high = 667 pM (100 ng/mL). **C** Incucyte® live cell imaging data from BRiDGE CAR or CD33 CAR T cells repeatedly encountering fresh cancer cells and AM. Left: U937 cell counts over time, new cancer cells added at each peak. Right: fold-change of BRiDGE CAR and CD33 CAR over time. N=2 donors, data presented as mean with SD. *P*-values calculated by two-way ANOVA with Šídák’s multiple comparisons test. **D** Memory subsets of BRiDGE CAR and CD33 T cells repeatedly exposed to fresh cancer cells and AM. Terminally differentiated effector memory T cells (T_EMRA_: CD45RA+CCR7−), effector memory (T_EM_: CD45RA−CCR7−), central memory (T_CM_: CD45RA−CCR7+) and naive (T_Naive_: CD45RA^+^CCR7^+^) T cell subsets presented as fraction of total CAR T cells. CD33 CAR T cells did not recover after fourth challenge. N=3 donors, data presented as mean. **E** *In vivo* pharmacokinetics study of CD33 AM. NSG mice were injected with indicated doses of CD33 AM. Plasma concentration of AM was measured by ELISA and half-life calculated independently per dose group, N=5 mice per group.

Repeated antigen challenge was used to mimic long-term engagement of BRiDGE CAR T cell function benchmarking it to a conventional CD33 CAR with the same signalling architecture, differing primarily in the mode of antigen engagement. Both systems controlled tumour growth without statistically significant differences, although BRiDGE CAR T cells showed a trend towards improved tumour control following repeated antigen challenges (Figure 3C, left). While overall expansion was comparable between the two groups, their expansion kinetics differed. Conventional CD33 CAR T cells underwent rapid early expansion followed by contraction, whereas BRiDGE CAR T cells exhibited more modest initial expansion but greater retention at later challenges (Figure 3C, right). Phenotypic analysis further revealed that BRiDGE CAR T cells retained higher proportions of naïve and central memory T cells, whereas conventional CD33 CAR T cells showed a greater shift towards effector memory and TEMRA phenotypes (Figure 3D). A similar trend was observed in a CD276 rechallenge model, in which BRiDGE CAR T cells maintained a higher proportion of central memory T cells than direct CD276 CAR T cells at later challenges (Supplementary Figure S3B). These results indicate that adaptor-mediated antigen engagement is associated with a less-differentiated phenotype and a larger residual CAR T cell population facing repeated antigen challenge.

We also investigated the pharmacokinetics of the CD33 AM to determine its *in vivo* persistence and potential dosing frequency. Following a single dose, plasma exposure was dose-dependent, with terminal half-lives of approximately 168, 300 and 258 h at 0.2, 1 and 5 mg/kg, respectively (Figure 3E). Half-lives at 1 and 5 mg/kg were comparable. These values align with the pharmacokinetics of conventional IgG antibodies and support periodic systemic dosing.

### *In vivo* tumour models demonstrate flexibility and control of BRiDGE CAR platform

*In vivo* efficacy and adaptor-mediated control of BRiDGE CAR T cells were further demonstrated across multiple tumour models. In the MOLM-13 (AML) xenograft model, BRiDGE CAR T cells with CD33 AM given twice weekly achieved sustained tumour control and significantly prolonged survival compared to BRiDGE CAR alone or conventional CD33 CAR T cells (Figure 4A). At study termination, CAR^+^ T cells were detected in blood and bone marrow of mice receiving BRiDGE CAR and AM, with minimal residual tumour cells (Figure 4B). IHC confirmed human T cell infiltration into liver and spleen (Supplementary Figure S4A). In a second model treating CD33^+^ U937 (AML), tumour control and survival were comparable between BRiDGE CAR with CD33 AM and conventional CD33 CAR T cells (Figure 4C). Both CAR T cell populations expanded and remained detectable during the study (Supplementary Figure S4B), while peak IL-2 and IFN-γ levels were lower with BRiDGE CAR than with conventional CD33 CAR T cells (Supplementary Figure S4C). In the CD19^+^ NALM-6 model (B-ALL), BRiDGE CAR T cells suppressed tumour growth and prolonged survival in an adaptor-dependent manner (Figure 4D).

**Figure 4.**
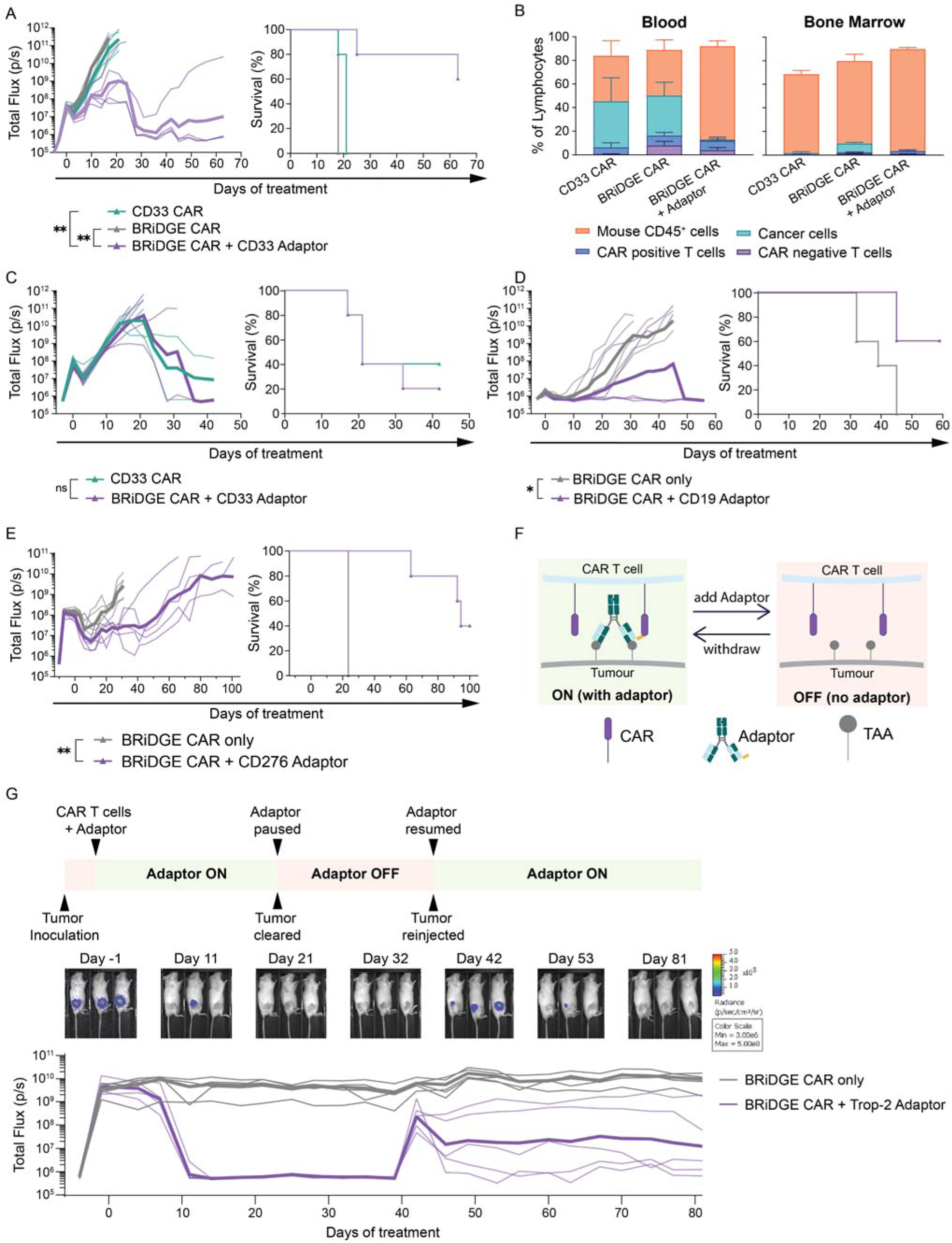
BRiDGE CAR efficacy treating *in vivo* models of leukaemia and solid cancers. All *in vivo* studies were designed with N=5 mice per group, 1 × 10□ target cells and 1 × 10□ CAR cells/animal. Treatment with CAR T cells began on day 0 with AMs administered i.p. twice weekly at a dose of 1.25 mg/kg. BLI plots show mean (thick line) and individual data points. *P*-values were calculated by log-rank test. **A** BLI and Kaplan–Meier survival analysis in a MOLM-13 model targeting CD33. The BRiDGE CAR only and BRiDGE CAR + Adaptor groups are the CAR only and XmAb monotag cohorts also shown in Figure 1F, G (*P*=0.0035 CD33 CAR vs BRiDGE CAR + Adaptor; *P*=0.0027 BRiDGE CAR only vs BRiDGE CAR + Adaptor). **B** Lymphocyte composition in blood (left) and bone marrow (right) of mice from A at study endpoint. **C** BLI summary and Kaplan–Meier survival analysis in a U937 model targeting CD33 (*P*=0.6568, CD33 CAR vs BRiDGE CAR + Adaptor). **D** BLI summary and Kaplan–Meier survival analysis in a NALM-6 model targeting CD19. In this study 0.5 × 10□ NALM-6 cells were engrafted (*P*=0.0173 BRiDGE CAR only vs BRiDGE CAR + Adaptor). **E** BLI summary and Kaplan–Meier survival analysis in a PC3 solid tumour model targeting CD276 (*P*=0.0027). **F** Mechanism of adaptor-mediated CAR activity. Without adaptor, CAR T cells cannot engage the antigen, and the cell remains inactive (off); adding the adaptor bridges the gap between CAR and target cell, activating the cell (on). Activity can be paused and resumed by withdrawing and re-applying the AM. **G** Experimental proof for mechanism presented in F. A localised solid tumour model of OVCAR-3 targeting Trop2^+^ was treated with BRiDGE CAR T cells and AM, once tumours had cleared for approximately one week (measured by BLI), AM was paused and after 20 days the same dose of cancer cells was injected at the same site and AM was resumed.

We next evaluated the BRiDGE CAR in solid tumour models targeting CD276 in PC3 (prostate cancer) and Trop2 in OVCAR-3 (ovarian cancer). In the CD276 model, BRiDGE CAR T cells with AM delayed tumour growth and significantly prolonged survival compared to BRiDGE CAR T cells alone (Figure 4E). For the Trop2 model, we mimicked a localised solid tumour mass by injecting cancer cells subcutaneously into the right flank. BRiDGE CAR T cells with AM reduced tumour signal to background levels in all mice by approximately day 11 (Figure 4G). A core premise of adaptor CAR systems is the ability to regulate CAR activation through AM dosing (Figure 4F). To test this control directly, we assessed whether BRiDGE CAR T cells could persist in the absence of AM and re-engage target cells once treatment with AM was readministered. Following tumour clearance, Trop2 AM was paused for 20 days, during which tumour signal remained at baseline. Mice were then rechallenged with another dose of OVCAR-3 cells and AM injection resumed without additional CAR T cell infusion (Figure 4G). Re-engagement of persisting BRiDGE CAR T cells was evident in three of five mice, in which tumour signal declined by approximately two logs and remained suppressed for the remainder of the study. CAR DNA was detected in the blood throughout the study even with adaptor withdrawal (Supplementary Figure S4D). These findings demonstrate adaptor-dependent tumour control across multiple tumour entities and the capability to re-engage persisting BRiDGE CAR T cells through adaptor re-dosing.

### Antigen density regulates BRiDGE CAR activity, with preserved haematopoietic progenitor function in vitro

As CD33 and CLL-1 are expressed on both AML and normal haematopoietic cells, differences in antigen density may influence BRiDGE CAR activity, with higher CD33 expression reported on AML than healthy haematopoietic stem cells [25]. We therefore examined how antigen density affects BRiDGE CAR activity. U937 single-cell clones spanning a range of CD33 or CLL-1 densities were selected by flow cytometry after CRISPR-Cas9 editing (Figure 5A, D). Increasing CD33 density reduced the AM concentration required for cytotoxicity, with cytotoxicity EC_50_ inversely related to antigen density (Figure 5B, C). IL-2 and IFN-γ EC_50_ values followed the same trend (Figure 5C; Supplementary Figure S5A). For CLL-1, cytotoxicity showed the clearest density dependence, with weaker correlations for IFN-γ and IL-2 secretion (Figure 5E, F; Supplementary Figure S5B). Together, these data show that BRiDGE CAR activity is dependent on both antigen density and AM concentration. CD33^+^ healthy PBMCs expressed approximately 1,000 CD33 molecules per cell (Figure 5G), well below expression measured on the U937 clones (4,464–24,123 molecules per cell).

**Figure 5.**
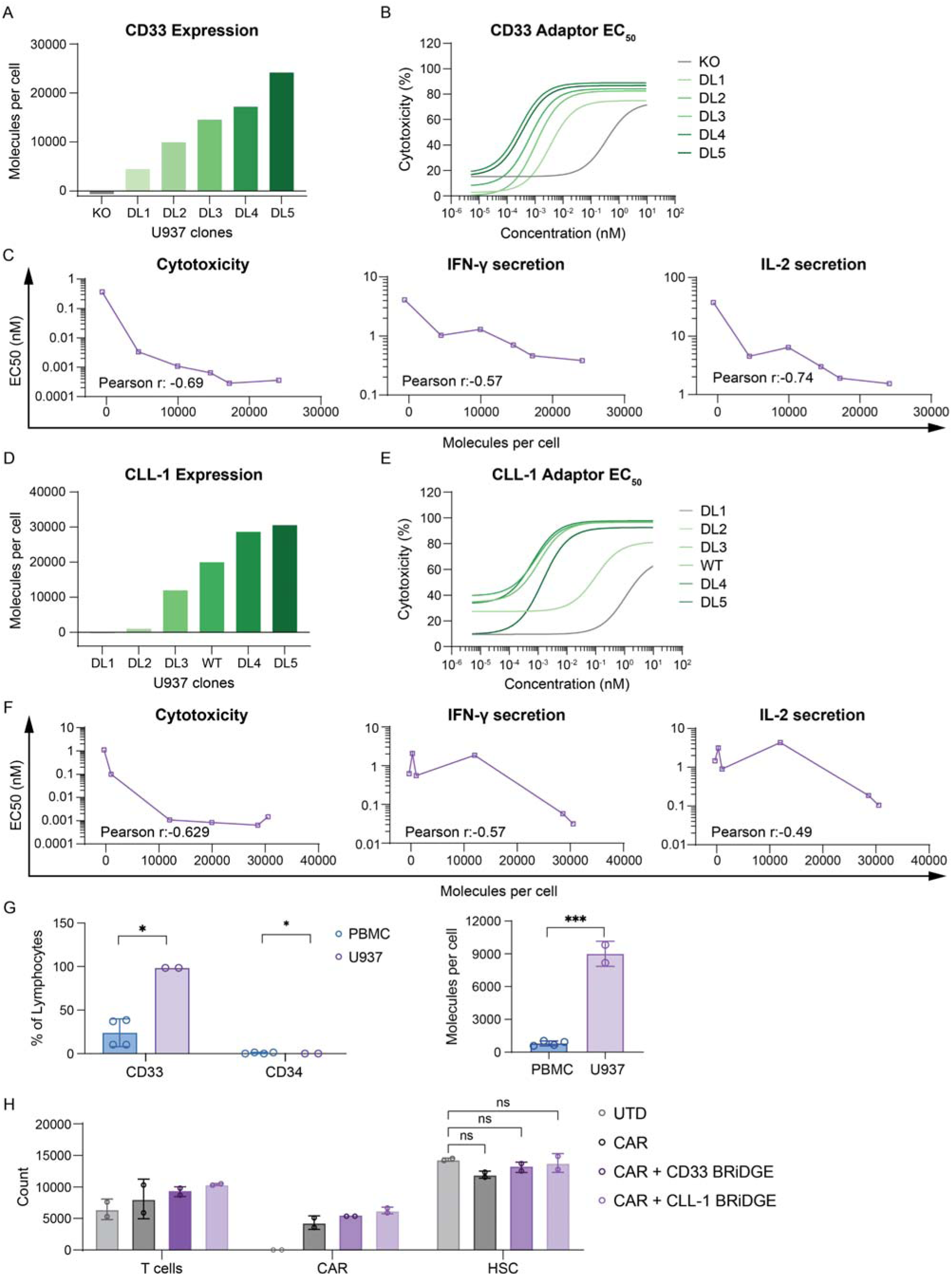
Activation of BRiDGE CAR is driven by antigen-density and preserves HSC function in vitro. **A** Quantification of CD33 molecules on U937 clones using QIFIKIT®. U937 CD33 KO: below the limit of detection, DL1: 4,464, DL2: 9,934, DL3: 14,519, DL4: 17,170 and DL5: 24,123; U937 WT was quantified (16,756) but not plotted. N=3 experiments, data presented as mean. **B** Non-linear regression curves calculated from BRiDGE CAR cytotoxicity with CD33 AM against U937 clones from A. **C** Correlation between CD33 density and EC_50_ for cytotoxicity (left), IFN-γ secretion (middle) and IL-2 secretion (right). Pearson r indicated on each graph. **D** Quantification of CLL-1 molecules on U937 clones using QIFIKIT®. U937 DL1: below the limit of detection, DL2: 980, DL3: 11,967, WT: 19,919, DL4: 28,593 and DL5: 30,509; U937 CLL-1 KO clone was quantified (264) but not plotted. N=1 experiment, data presented as mean. **E** Non-linear regression curves calculated from BRiDGE CAR cytotoxicity with CLL-1 AM against U937 clones from D. **F** Correlation between CLL-1 density and EC_50_ for cytotoxicity (left), IFN-γ secretion (middle) and IL-2 secretion (right). Pearson r indicated on each graph. Non-linear regression curves and EC_50_ were calculated using a three-parameter agonist vs. response nonlinear regression model. **G** CD33 and CD34 expression on human PBMC from healthy donors and on parental U937 cells. Left: CD33□ and CD34□ cells as a percentage of live cells. Right: quantification of CD33 molecules per cell on the same populations using QIFIKIT®. N=2 donors with 2 technical replicates each, data presented as mean with SD. **H** Haematopoietic stem cell (HSC), CAR T cell and total T cell counts after 4 h of co-incubation. *P*-values calculated by two-way ANOVA with Tukey’s multiple comparisons test (P=0.1102 UTD vs CAR; P=0.3644 UTD vs CAR + CD33 BRiDGE; P=0.9179 UTD vs CAR + CLL-1 BRiDGE).

We next assessed potential toxicity against haematopoietic stem cells (HSCs). Colony formation was unchanged after 14 days of culture with BRiDGE CAR T cells, alone or with CD33 or CLL-1 AM (Supplementary Figure S5C), and HSC numbers were likewise unaffected in a 4 h co-culture (Figure 5H). HSCs showed minimal Bil03 binding and CD33 expression, whereas CLL-1 was expressed on approximately 19% of HSCs (Supplementary Figure S5D). That colony formation was preserved in the presence of the CLL-1 AM suggests this level of expression remains below the threshold required for BRiDGE CAR activation.

### IL-18 armoured BRiDGE CAR shows increased potency *in vitro* and *in vivo*

Interest in IL-18 armouring as a strategy to enhance CAR T cell activity has continued to grow in recent years [26, 27]. To assess whether the BRiDGE platform could incorporate an inducible cytokine module, we placed human IL-18 under the T cell-specific iSynpro promoter [28] downstream of the BRiDGE CAR sequence (Figure 6A). The addition of the inducible IL-18 module did not impair CAR transduction, expansion, memory phenotype or expression of PD-1, LAG-3 and TIM-3 (Figure 6B; Supplementary Figure S6B–D). There was comparable cytotoxicity of BRiDGE CAR-IL18 and non-armoured BRiDGE CAR T cells in the CD33 model targeting U937 cells across the tested dose range (Figure 6C), whereas BRiDGE CAR-IL18 exerted significantly improved cytotoxicity in the CD276 model, especially at lower AM concentrations (Supplementary Figure S6E).

**Figure 6.**
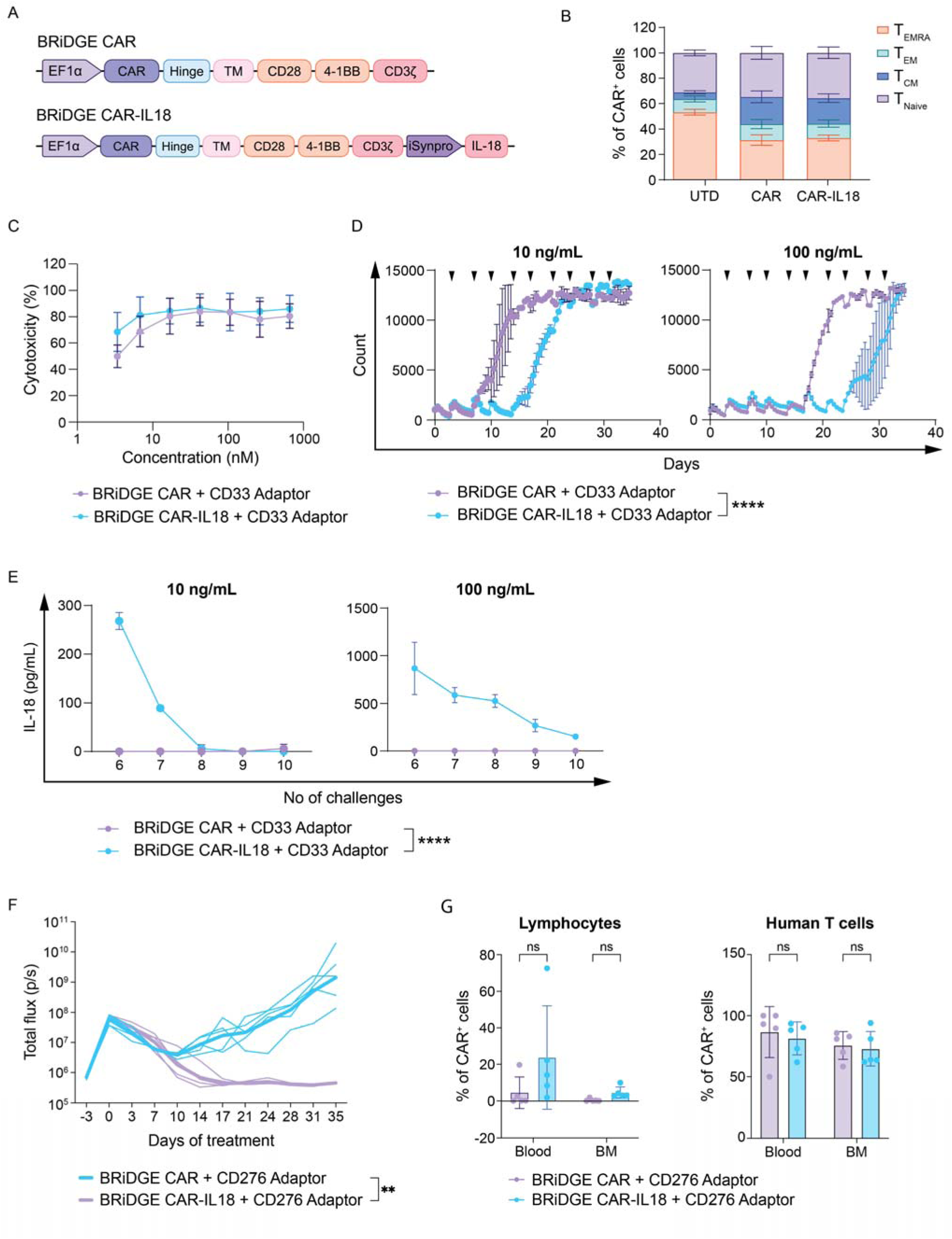
Activity of BRiDGE CAR T cells armoured with IL-18 *in vitro* and *in vivo*. **A** Schematic of the BRiDGE CAR and BRiDGE CAR-IL18 constructs. **B** Memory subsets (T_EMRA_, T_EM_, TCM and Tnaive) of UTD, BRiDGE CAR and BRiDGE CAR-IL18 T cells 5 days after transduction. N=4 donors, data presented as mean with SD. **C** CD33 AM mediated cytotoxicity against U937 by armoured vs non-armoured BRiDGE CAR T cells after 24 h. N=3 donors, data presented as mean with SD. **D** U937 cell counts over time during a rechallenge assay of armoured CAR T cells with 10 ng/mL (67 pM; left) and 100 ng/mL (667 pM; right) CD33 AM, addition of fresh cancer cells and AM indicated by arrows. Representative of two donors with consistent relative differences between constructs; data from the representative donor presented as mean with SD of technical replicates. *P*-values calculated by two-way ANOVA with Tukey’s multiple comparisons test (*P*<0.0001 BRiDGE CAR vs BRiDGE CAR-IL18). **E** ELISA detecting IL-18 secretion in supernatant from D, challenges 6–10 tested. N=2 donors, data presented as mean with SD. *P*-values calculated by two-way ANOVA with Tukey’s multiple comparisons test (*P*<0.0001 BRiDGE CAR vs BRiDGE CAR-IL18). **F** Bioluminescence imaging of a CD276□ prostate cancer mouse model. Treatment with 1 × 10□ CAR□ cells/animal, and CD276 AM dose as stated in Figure 4. Data presented as mean with SD, *P*-values calculated by Mixed-effects model (REML) with Tukey’s multiple comparisons test (*P*=0.0027 on day 35). **G** Quantification of CAR^+^ cells in blood and bone marrow (BM) from animals in F at study endpoint relative to the lymphocyte (left) and human T cell (right) subset, data presented as mean with SD, *P*-values calculated by unpaired t test with Welch’s correction (Lymphocytes: *P*=0.207243 in blood and *P*=0.073957 in BM; T cells: *P*=0.881956 in blood and BM).

Upon repeated antigen challenge with U937 cells, BRiDGE CAR-IL18 T cells delayed tumour cell growth by approximately one week at both 10 and 100 ng/mL CD33 AM (Figure 6D). IL-18 secretion declined over successive challenges, reaching baseline by the eighth rechallenge at 10 ng/mL while remaining detectable at 100 ng/mL until the end of the experiment (Figure 6E).

Finally, we evaluated IL-18 armouring *in vivo* using a disseminated CD276□ PC3 model at one-tenth of the CAR T-cell dose used in our earlier experiments (1 × 10□ versus 1 × 10□ CAR□ cells per animal). Both armoured and non-armoured BRiDGE CAR T cells initially reduced tumour burden. However, tumours subsequently progressed in mice receiving non-armoured CAR T cells, whereas BRiDGE CAR-IL18 T cells maintained tumour control throughout the study (Figure 6F). At study endpoint, armoured recipients showed a non-significant trend toward a higher CAR□ proportion of lymphocytes in blood and bone marrow, with comparable CAR□ fractions within the human T cell compartment (Figure 6G). As expected, IL-18 was detected only in serum from armoured CAR T cells (Supplementary Figure S6F). These findings demonstrate that inducible IL-18 armouring enhances BRiDGE CAR T-cell activity during repeated antigen challenge and improves tumour control *in vivo* under reduced CAR T-cell dosing conditions.

## Discussion

The BRiDGE CAR platform separates antigen recognition from CAR signalling, enabling systematic evaluation of the design parameters that govern adaptor-mediated CAR T-cell activity. We identified key features of CAR and adaptor design that influence antitumour efficacy and demonstrated that a single CAR construct can mediate tumour clearance across multiple haematological and solid tumour models through different targeting adaptors. Compared with conventional direct-targeting CAR T cells, BRiDGE CAR T cells also exhibited distinct expansion kinetics and greater retention of less-differentiated phenotypes during repeated antigen challenge, suggesting that adaptor-mediated antigen engagement may influence CAR T-cell responses beyond its role in directing target recognition.

Productive adaptor-mediated recognition depends on the configuration of both the CAR and the adaptor. Consistent with previous studies showing that CAR geometry influences activity [29–31], BRiDGE CAR function varied with hinge length and adaptor architecture. Full-length bivalent adaptors showed greater potency than Fab-based or monovalent formats, indicating contributions from both antibody format and binding valency. In contrast, increasing E200 tag valency provided little additional benefit *in vitro* and was associated with less durable tumour control *in vivo*. One possibility is that multiple CAR binding tags promote receptor cross-linking or bridging between neighbouring CAR T cells, a mechanism proposed for adaptor systems [20] but not yet demonstrated experimentally. Together, these findings suggest that effective modular CAR design depends on the geometry of the CAR–adaptor–antigen interaction and on the distinct features of target and CAR binding, rather than simply maximising binding valency.

In vivo, adaptor-mediated recognition achieved at least equivalent activity to direct CD33 CAR recognition in both models tested, matching it in U937 and exceeding it in MOLM-13. The mode of antigen engagement may also influence how CAR T cells respond to repeated stimulation. Compared with direct CD33 CAR T cells using the same signalling architecture, BRiDGE CAR T cells in conjunction with the CD33 AM showed more modest early expansion, greater persistence and better preservation of less-differentiated phenotypes during repeated antigen challenge. A similar preservation of central memory cells was observed with CD276 targeting, suggesting that this behaviour was not restricted to a single antigen. Recent studies have shown that adaptor concentration can alter CAR T cell activation states and metabolic programmes [32], while treatment-free intervals can mitigate chronic stimulation and T cell dysfunction in adaptor CAR systems [33]. Our findings complement these observations by showing distinctive responses over time, separating adaptor-mediated and direct antigen recognition during repeated challenge. Whether these differences arise from reversible CAR–AM binding, reduced cumulative signalling or other features of adaptor-mediated interaction will require further investigation.

Adaptor dosing provides a means for controlling CAR T cell activity. The response depended on both AM concentration and antigen density, with higher antigen density requiring less adaptor to achieve cytotoxicity. This relationship may be particularly relevant in AML, where candidate targets such as CD33 are shared between malignant and normal haematopoietic cells and selectivity cannot rely solely on antigen presence or absence [25]. Adaptor dosing also provides temporal control. BRiDGE CAR T cells persisted during adaptor withdrawal and could subsequently be re-engaged by resuming AM treatment without an additional CAR T cell dose. A full-length IgG adaptor has a long half-life, which permits infrequent dosing but means activity cannot be switched off quickly; in our rechallenge experiment the adaptor was withheld for 20 days before tumour cells were re-injected. Thus, the adaptor functions not simply as a targeting bridge, but also as a pharmacological tool for controlling CAR T cell engagement.

The E200–Bil03 interaction enables targeting and cellular function to be modified independently. A CD33×CLL-1 bispecific adaptor enabled the same BRiDGE CAR T cells to recognise either AML antigen, illustrating how adaptor design could address antigen heterogeneity without generating a new cellular product. Clinically established dual-targeting antibody formats, including amivantamab (EGFR×MET) [34] and zanidatamab (biparatopic HER2) [35], provide potential frameworks for developing such adaptors. Our bispecific results, however, are currently limited to *in vitro* activity and require *in vivo* validation.

IL-18 armouring controlled by an inducible promoter enhanced activity during repeated antigen challenge and improved tumour control at a reduced CAR T cell dose. IL-18-armoured CAR T cells have shown enhanced activity in preclinical models [36–38] and, more recently, clinical activity in patients with B-cell lymphoma, even after previous CAR T cell therapy [27]. BRiDGE CAR applies this approach to a modular CAR architecture, showing that inducible cytokine enhancement can be combined with adaptor-mediated antigen recognition.

Our study has several limitations that guide the next steps of the development of the BRiDGE CAR technology. The *in vivo* studies were performed in immunodeficient NSG mice, limiting assessment of host immune interactions, and particularly the broader effects of IL-18 armouring. The relationship between adaptor dose, antigen density and healthy tissue targeting needs to be confirmed *in vivo*, additionally bispecific CD33×CLL-1 targeting has only been studied *in vitro*. Residual killing of antigen-negative targets at high adaptor concentrations is consistent with allogeneic recognition of tumour cell lines by donor-derived T cells, and specificity is therefore most reliably assessed in the mixed co-culture format, where only antigen-positive populations were depleted (Figure 2D). Another important topic is the tissue cross-reactivity assessment of the E200 epitope, which is currently ongoing. Addressing these questions in more physiologically relevant models will be important for clinical translation.

In summary, our BRiDGE CAR technology demonstrates that separating antigen recognition from CAR signalling can support antitumour activity whilst providing external control over antigen engagement. Our study defines design principles governing adaptor-mediated recognition and shows that adaptor concentration, antigen density and the mode of antigen engagement each shape CAR T cell activity. The ability to independently modify targeting and cellular function supports the broader application of this modular architecture across different tumour settings.

## Materials and methods

### Generation of Lentiviral Vectors

Third-generation lentiviral vectors (LV) were produced by transfecting Lenti-X cells (TakaraBio) using Lipofectamine 3000 (Thermo Fisher) according to the manufacturer’s protocol. Supernatant containing lentiviral vectors was harvested twice, at 24 h and 48 h post-transfection. Harvests were clarified, filtered and concentrated 2 days post-transfection. LV pellets were incubated on ice for 1 h and resuspended in either Tris-HCl (pH 8.0) or Tris-NaCl buffer (25 mM Tris, 60 mM NaCl, pH 7.5), depending on the preparation. Concentrated lentiviral vectors were aliquoted and stored at −80°C.

### Protein Expression and purification

Recombinant antibodies were transiently expressed in ExpiCHO-S cells using ExpiFectamine CHO transfection reagents according to the manufacturer’s protocol (Thermo Fisher). Cells were cultured in ExpiCHO Expression Medium, and supernatant was harvested 7 days post-transfection and clarified by centrifugation, filtered and purified. Samples were purified using the ÄKTA pure chromatography system (Cytiva). Proteins were captured using protein A affinity chromatography, followed by cation exchange chromatography. Protein quality and purity were assessed by SDS-PAGE and analytical size-exclusion chromatography (SEC).

### Cell lines and culturing

All cell lines were acquired from ATCC: U937, MOLM-13, NALM-6, THP-1, AML-193, HL-60, JeKo-1, OVCAR-3 and PC3. They were maintained in complete RPMI: RPMI 1640 Medium, GlutaMAX™ Supplement (Thermo Fisher) supplemented with 10% heat-inactivated FBS and 1% Penicillin-Streptomycin (10,000 U/mL, Thermo Fisher). JeKo-1 (CD19□CD33□) was used both as a CD19 target model and, in CD33-directed experiments, as a CD33-negative control.

### Generation of luciferase-expressing cell lines

Transfer plasmids encoding luciferase and eGFP were generated in-house by modifying the third-generation lentiviral vector (Addgene #12252). Lentiviral particles were produced as described above. Cancer cell lines were transduced and enriched accordingly. Flow cytometry was used to confirm transgene expression.

### Flow cytometry

Samples were stained in FACS buffer containing Dulbecco’s phosphate buffered saline (DPBS; Thermo Fisher) with 1% FBS for 10 min at room temperature, shielded from light. Data were acquired using a MACSQuant® 16 and analysed with FlowJo™. AM binding was validated by a two-step staining protocol: cell lines were incubated with AMs, followed by a secondary antibody (Bil03) conjugated to AF647 that recognises the E200-tag. All antibodies, i.e. Biotin-VioBlue (Bio3-18E7), CD4-PE-Vio615 (REA623), CD8-APC-Vio770 (REA734), CD25-PE (REA570), CD45RA-PerCP-Vio700 (REA562), CD69-APC (REA824), CCR7-FITC (REA546), LAG3-APC (REA351), PD-1-PE (PD1.3.1.3) and TIM3-PE-Vio770 (REA635) were used in accordance with manufacturer’s instructions (Miltenyi Biotec).

### PBMC isolation and generation of CAR T cells

Peripheral blood from healthy donors was acquired from the Australian Red Cross Lifeblood®. Peripheral blood mononuclear cells (PBMCs) were isolated using density gradient separation with Ficoll-Paque PREMIUM (Cytiva). CD4^+^ and CD8^+^ T cells were isolated separately using MACS® microbeads and magnetic cell separators (Miltenyi Biotec).

For CAR T generation, CD4^+^ and CD8^+^ T cells were seeded at a 1:1 ratio in TexMACS™ supplemented with IL-7, IL-15 (10/15 ng/mL, Miltenyi Biotec) and FBS (3%, Thermo Fisher), and activated with TransAct™ (Miltenyi Biotec). 48 h later, lentiviral particles were added. Transduction was confirmed after 7 days using a biotinylated CAR detection reagent (generated in-house), followed by staining with Biotin-VioBlue. CAR T cells were expanded for 14 days and then cryopreserved in FBS with 10% DMSO.

### Luciferase-based cytotoxicity assay

Tumour cells, AM and effector cells were plated in 96-well plates (Costar) with 1 µg/mL of synthetic D-Luciferin (Sigma-Aldrich). Plates were incubated at 37°C and 5% CO_2_ for up to 72 h. SpectraMax® i3 (Molecular Devices) was used to measure the plates at defined timepoints. Cytotoxicity was calculated using the relative luminescence of testing conditions compared to standard controls. Effector-to-target ratios, AM concentration ranges and incubation times differed between experiments and are specified in the corresponding figure legends.

### Cytokine secretion assay

Tumour cells, AMs and effector cells were plated in 96-well plates (Corning). Plates were incubated at 37°C and 5% CO_2_ for 24 h. Supernatant was harvested after centrifugation and stored at −20°C until analysis. Human MACSPlex® Cytokine 12 kit (Miltenyi Biotec) was used to measure cytokine concentration following the manufacturer’s instructions. Samples were read on MACSQuant® 16 and analysed using FlowJo™.

### Flow cytometry-based *in vitro* cytotoxicity assay

Tumour cells were incubated with CAR T cells and AM at 37°C and 5% CO_2_ for 24 h, washed and stained with fluorescent antibodies as mentioned above. Samples were read on a MACSQuant® 16 and data analysed using FlowJo™.

### Generation of antigen knockout variants using CRISPR/Cas9 technology

CD33 and CLL-1 Knockout (KO) variants were generated from our eGFP/Luc expressing reporter cell line, U937. To prepare gRNA, crRNA (IDT) and tracrRNA (IDT) were mixed, denatured in a PCR thermocycler (Thermo Fisher), followed by annealing and gradual cooling to RT. The ribonucleoprotein complex (RNP) was prepared by incubating the gRNA and Cas9 for 10 min at room temperature. SF-nucleofection kit and Amaxa 4D nucleofector (both Lonza) were used for electroporation according to manufacturer’s instructions. KO efficiency was tested using flow cytometry, followed by single-cell sorting to generate stable clones.

### Incucyte live cell imaging

GFP-expressing U937 cells, AM and CAR T cells were plated in 96-well plates. Cells were kept in an incubator containing the Incucyte S3 system (Sartorius) and cultured for 3 to 4 weeks at 37°C and 5% CO_2_. Twice weekly, fresh U937, AM and medium were added to CAR T cells. Fluorescent images were acquired every 12 h. Sartorius Incucyte software was used for readout and data analysis.

### IL-18 ELISA

Cell culture supernatant was diluted 1:10 and mouse plasma was diluted 1:100, 1:500, 1:1000 & 1:2000. IL-18 concentration was measured using the Human Total IL-18 DuoSet ELISA (R&D Systems) according to the manufacturer’s instructions.

### Colony-forming unit (CFU) assay

Cord blood-derived CD34^+^ cells were purchased from Stem Cell Technologies and co-incubated with CAR T cells at a 10:1 ratio for 4 h. Cells were then transferred into 35 mm dishes with MethoCult™ H4034 Optimum medium containing rhSCF, rhIL-3, rhEPO, rhG-CSF and rhGM-CSF (Stem Cell Technologies). Colonies were counted after 14 days of culture.

### Digital PCR

CAR gene copy number was measured using the QIAcuity digital PCR (dPCR) system. Briefly, DNA was isolated using the GeneJET Genomic DNA purification kit (Thermo Fisher) and mixed with the PCR Master Mix (QIAcuity Probe PCR Kit, Cat# 250101), probes detecting the RRE (Bio-Rad, Cat# 10042958, Assay ID: dCNS153736898) and RPP30 (Cat# 10031244, Assay ID: dHsaCP1000485) region, digestion enzymes and buffers in a 24 nanoplate (QIAcuity Nanoplate 26k 24-well, Cat# 250031) and amplified in the QIAcuity instrument (40 cycles) before imaging.

### *In vivo* experiments

6- to 8-week-old female NOD.Cg-Prkdcscid Il2rgtm1Wjl/SzJ (NSG) mice were sourced from Australian BioResources (ABR). Cancer cells were injected into the tail vein for metastatic models or the right flank for subcutaneous models 3–7 days prior to treatment. On day 0, all animals underwent bioluminescence imaging (BLI) using the IVIS Lumina X5 to determine baseline values. CAR T cells were then injected intravenously, and AM dosing commenced on the same day at 1.25 mg/kg (equivalent to 25 µg per animal, i.p., twice weekly). Tumour growth was monitored twice weekly by BLI. After study termination, blood and bone marrow samples were harvested and stained using CD3, msCD45, Biotin-VioBlue and our detection reagent using the method described above. Other organs of interest were formalin-fixed and paraffin-embedded for immunohistochemical analysis at the Westmead Institute for Medical Research. Sections were stained for human CD3 using a polyclonal rabbit anti-human CD3 primary antibody (Dako, A0452), followed by HRP-conjugated goat anti-rabbit immunoglobulins as secondary antibody (Dako, P0448).

### Software

Flow cytometry data were analysed using FlowJo™ v10.10. Living Image® v4.8.4 was used for BLI acquisition and analysis. Incucyte® Software v2024A was used for Incucyte live cell imaging data acquisition and analysis. SoftMax® Pro v7.1 was used for SpectraMax® data acquisition. NDP.view 2.9.29 was used for analysis of immunohistological slides. QIAcuity Software v3.2 was used for dPCR analysis. Graphs and statistical analyses were performed in GraphPad Prism v10.1.

### Statistics

GraphPad Prism v10.1 was used for plotting and statistical analyses. Statistical tests applied are specified in the corresponding figure legends. Statistical significance was defined as *P* < 0.05.

## Acknowledgements

Preclinical imaging was performed at the Westmead Scientific Platforms, which are supported by the Westmead Research Hub, the Cancer Institute New South Wales, the National Health and Medical Research Council and the Ian Potter Foundation. We thank D. Christ, J. Henry and H. Lenthal for affinity testing and discussions.

## Author contributions

Conceptualisation: Z.L. Methodology and experimental planning: Z.L., A.H.J., M.M. Investigation: A.H.J., M.M., R.M., T.K., E.K., J.Y.; vector and adaptor production by E.K. and J.Y.; protein purification by R.M. and M.Y. Formal analysis: A.H.J., M.M., R.M., T.K. Visualisation: A.H.J., M.M., R.M., T.K. Data interpretation: Z.L., A.H.J., M.M., D.B. Writing – original draft: Z.L. and A.H.J. Writing – review and editing: Z.L., A.H.J., M.M., D.B. and all authors. All authors reviewed and approved the final manuscript.

## Ethics statement

In vivo study protocols were reviewed and approved by the Westmead Hospital Animal Ethics Committee and were performed in compliance with Australian animal welfare legislation and institutional policies. Human ethics approval was obtained from the Sydney Children’s Hospital Ethics Committee.

## Data availability

The data supporting the findings of this study are available within the paper and its Supplementary Information. Additional data, including source data underlying the figures, are available from the corresponding author on reasonable request.

## Code availability

No custom computer code was generated in this study. Data were analysed using commercially available software: FlowJo v10.10, GraphPad Prism v10.1, Living Image v4.8.4, Incucyte Software v2024A, SoftMax Pro v7.1, NDP.view 2.9.29 and QIAcuity Software v3.2.

