## Supplementary Figures for "Development and design rules for a modular human peptide-tag adaptor CAR T cell system"

### Supplement

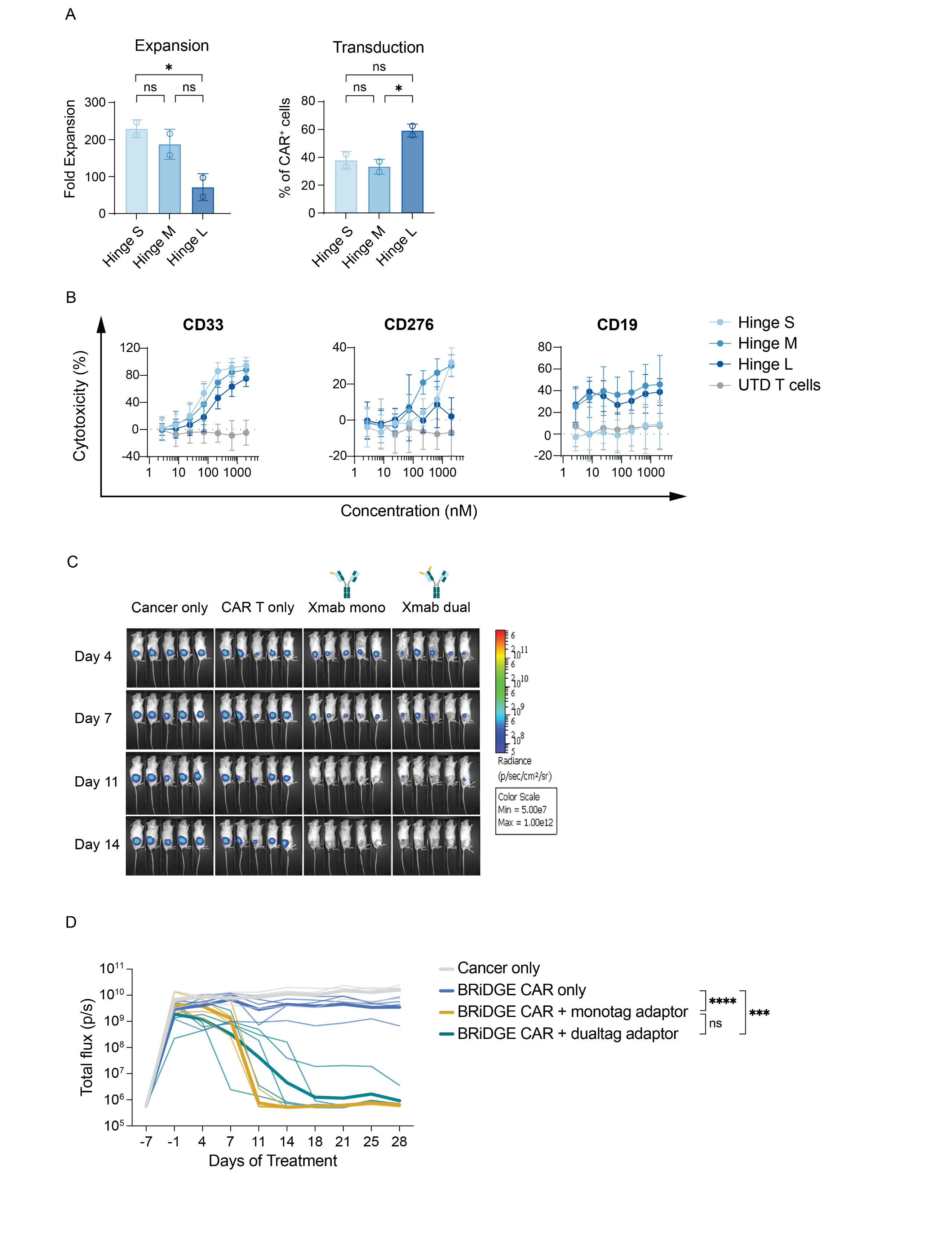

**Supplementary Figure S1 BRiDGE CAR characterisation and *in vivo* comparison of AM formats**

**A** Transduction and expansion of BRiDGE CAR constructs equipped with a short, medium or long hinge. Expansion is shown on the left (short vs long hinge *P*=0.0396) and transduction on the right (medium vs long hinge *P*=0.0358). Exemplary data from N=2 donors (independent CAR T cell production runs), five technical replicates per donor. Data presented as mean with SD, one-way ANOVA with Tukey’s multiple comparisons test. **B** Impact of short, medium or long hinge length on BRiDGE CAR cytotoxicity (N=2 donors, five technical replicates each) in a CD33 (MOLM-13), CD276 (PC3) and CD19 (JeKo-1) model, evaluated by luciferase assay; TAA indicated above graphs **C** BLI of OVCAR-3 bearing mice receiving treatment with Trop2 monotag or dualtag (asymmetrical) AM, N=5 mice per group. **D** Quantification of BLI signal from C, expressed as total flux (photons/s) over time, including pre-treatment baseline and timepoints not shown in C.

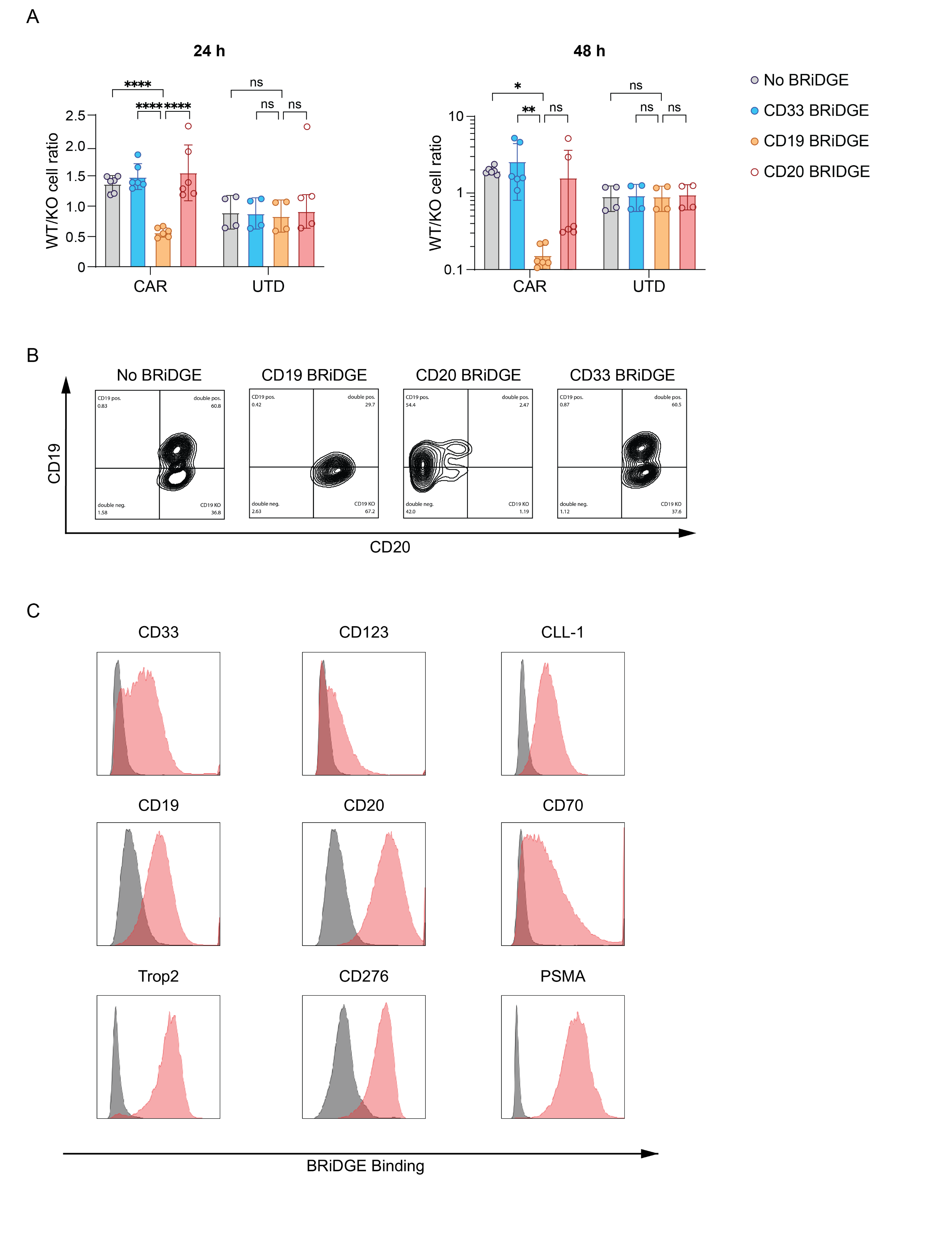

**Supplementary Figure S2 Specificity of BRiDGE CAR using CD19/CD20 AM and flow cytometric binding of AM**

BRiDGE CAR antigen specificity was assessed using a 1:1 mixture of CD19-KO and WT populations of JeKo-1 cells. Effector cells were co-cultured with target cells at an E/T ratio of 1:1 in the presence of CD19, CD20 or CD33 (neg. control) targeting AMs. 24 h and 48 h after treatment, tumour cells were stained for CD19 and CD20 expression. **A** Ratio of JeKo-1 WT to CD19-KO variants, N=3 donors, data presented as mean with SD. *P*-values calculated by two-way ANOVA with Tukey’s multiple comparisons test. *: *P*<0.05, **: *P*<0.01, ****: *P*<0.0001, ns: not significant**. B** Representative flow plots from A (one donor). **C** Representative plots of flow cytometric binding evaluation of nine BRiDGE AMs to antigen positive target cells. BRiDGE binding was detected using Bil03, shown in red, and Bil03 staining without AM is shown in grey.

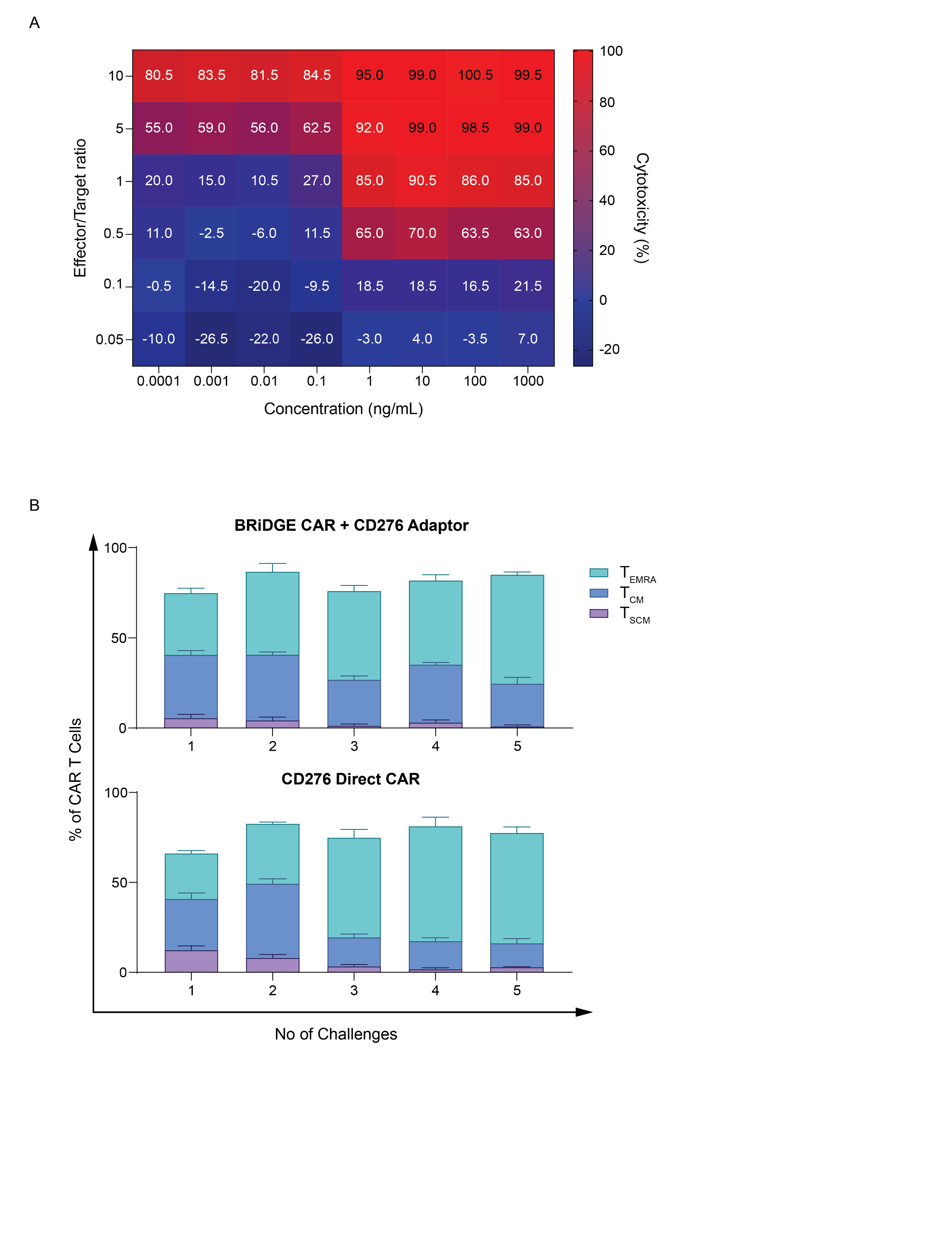

**Supplementary Figure S3 Dose-dependency of BRiDGE CAR cytotoxicity and memory phenotype over multiple challenges**

**A** Titration of effector-to-target (E/T) ratio and BRiDGE molecule concentration demonstrating dose-dependent cytotoxicity of BRiDGE CAR T cells. N=2 donors. **B** CAR T cell memory phenotype over multiple challenges with CD276^+^ PC3 cells. T_SCM_, T_CM_, and T_EMRA_ subsets are depicted as fraction of total CAR T cells. N=3 donors, data presented as mean with SD.

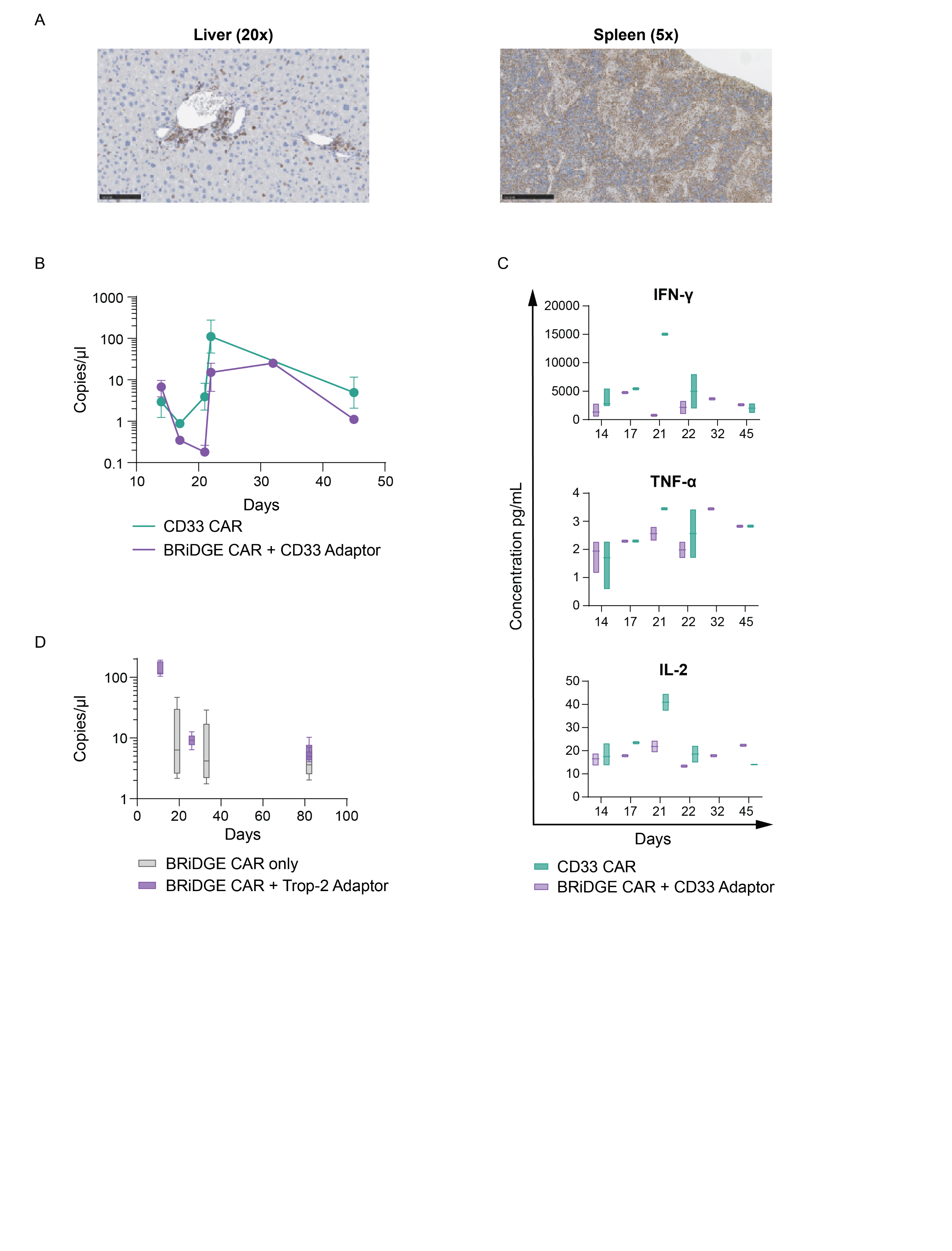

**Supplementary Figure S4 Tissue infiltration, expansion, and cytokine secretion of BRiDGE CAR T cells *in vivo***

**A** Immunohistochemistry staining for human CD3 on tissue sections collected 63 days after treatment from MOLM-13 bearing mice that received BRiDGE CAR T cells with CD33 AM. Representative images of liver (20× magnification, left), spleen (5× magnification, right); human CD3 stained in brown. **B** Quantification of CAR DNA in peripheral blood over time from the CD33^+^ U937 *in vivo* model. **C** Serum concentrations of IFN-γ, TNF-α and IL-2 over time in the groups from B. **D** Quantification of CAR DNA in peripheral blood over time from the Trop2^+^ OVCAR-3 *in vivo* model. Data presented as mean with SD and include animals remaining in each group at the specified timepoint (N=1–5).

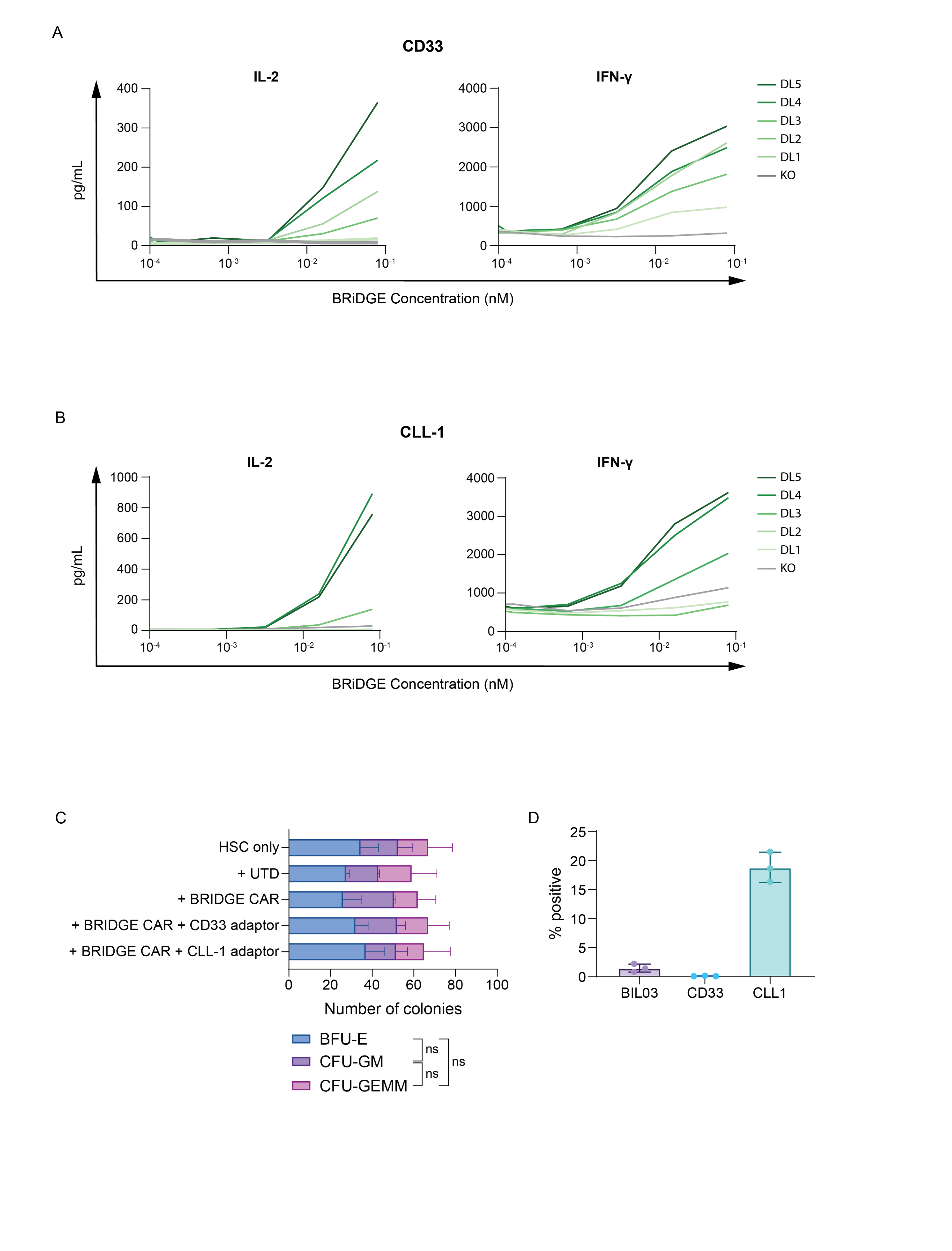

**Supplementary Figure S5 Antigen-density dependence of cytokine production and HSC toxicity *in vitro***

**A** IL-2 (left) and IFN-γ (right) secretion of BRiDGE CAR at six levels of CD33 expression including a CD33 KO control. **B** IL-2 (left) and IFN-γ (right) secretion of BRiDGE CAR at six levels of CLL-1 expression including a CLL-1 KO control. Antigen densities corresponding to each dose level are depicted in Figure 5A and 5D. N=2 donors, data presented as mean. **C** Colony-forming-unit (CFU) assay of HSC cultured alone or with untransduced (UTD) T cells, BRiDGE CAR T cells alone, or BRiDGE CAR T cells with either CD33 or CLL-1 AM for 14 days. N=2 donors, data presented as mean with SD. *P*-values calculated by two-way ANOVA with Tukey’s multiple comparisons test. BFU-E: Burst forming unit – erythroid; CFU-GM: Colony forming unit – Granulocyte, Macrophage; CFU-GEMM: Colony forming unit – Granulocyte, Erythrocyte, Monocyte, Megakaryocyte. **D** Surface staining of HSC with our E200 epitope targeting antibody Bil03 as well as CD33- and CLL-1-targeting antibodies. N=3 donors, data presented as mean with SD.

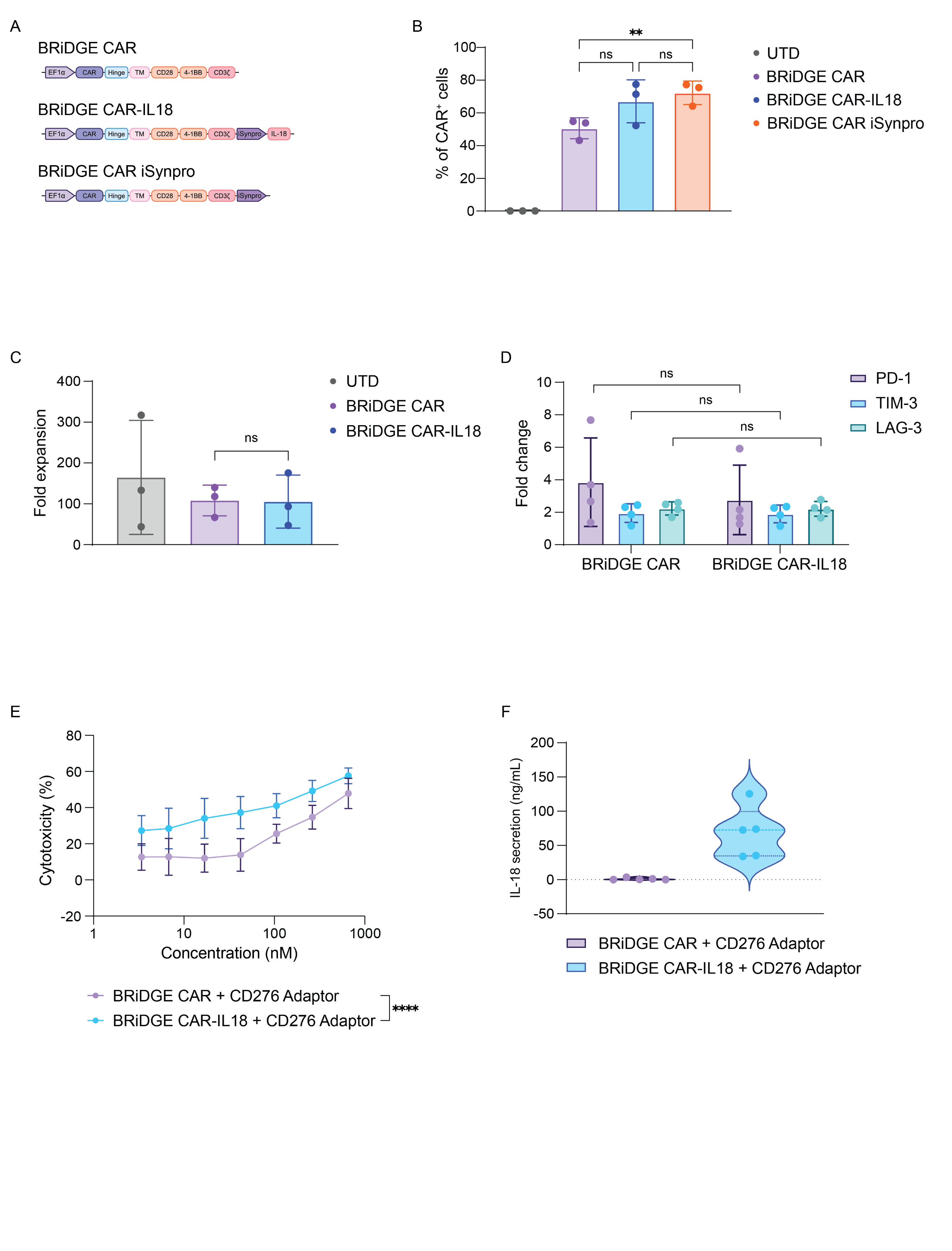

**Supplementary Figure S6 Further characterisation of BRiDGE CAR armoured with IL-18**

**A** Schematics of the BRiDGE CAR, BRiDGE CAR-IL18 and BRiDGE CAR iSynpro constructs (top to bottom). BRiDGE CAR-IL18 carries iSynpro-driven IL-18 and BRiDGE CAR iSynpro carries iSynpro alone to serve as a control. **B** CAR expression of armoured BRiDGE CAR T cells 5 days after transduction. BRiDGE CAR vs BRiDGE CAR iSynpro *P*=0.0036. **C** Expansion of armoured BRiDGE CAR constructs after 14 days plotted as fold expansion from baseline. B, C N=3 donors, data presented as mean with SD. *P*-values calculated by one-way ANOVA with Tukey’s multiple comparisons test. **D** Expression of PD-1, TIM-3 and LAG-3 in armoured BRiDGE CAR T cells 5 days after transduction. N=4 donors, data presented as mean with SD, plotted as fold increase from baseline. *P*-values calculated by two-way ANOVA with Šídák’s multiple comparisons test. **E** BRiDGE CAR and BRiDGE CAR-IL18 cytotoxicity with CD276 AM against PC3. N=3 donors, data presented as mean with SD. *P*-values calculated by two-way ANOVA with Šídák’s multiple comparisons test, *P*<0.0001. **F** IL-18 concentration in serum from BRiDGE CAR and BRiDGE CAR-IL18 treated mice, N=5 mice per group, data presented as violin plots with median and quartiles.
